# Proteasome autophagy is specifically regulated and requires factors dispensible for general autophagy

**DOI:** 10.1101/2021.03.26.437055

**Authors:** Kenrick A. Waite, Alicia Burris, Gabrielle Vontz, Angelica Lang, Jeroen Roelofs

## Abstract

Changing physiological conditions can increase the need for protein degradative capacity in eukaryotic cells. Both the ubiquitin-proteasome system and autophagy contribute to protein degradation. However, proteasomes are also an autophagy substrate. Thus, these processes must be differentially regulated depending on the physiological conditions presented. The signals and molecular mechanisms that govern proteasome autophagy are only partly elucidated. Our data indicate that chemical inhibition of TORC1 with rapamycin induces a bi-phasic response where proteasome levels are upregulated followed by an autophagy-dependent reduction. Surprisingly, several conditions that result in inhibited TORC1 exclusively induce proteasome autophagy (i.e. without any proteasome upregulation), suggesting a convergence of signals upstream of proteasome autophagy under different physiological conditions. Indeed, several conditions that activate general autophagy did not induce proteasome autophagy further distinguishing between proteasome autophagy and general autophagy. Consistent with this, we found that Atg11, the receptor for selective autophagy, and the map kinases Mpk1, Mkk1, and Mkk2, all play a role in autophagy of proteasomes, while they are dispensible for general autophagy. In all, our data provide new insights into the molecular regulation of proteasome autophagy by demonstrating that these complexes are specifically regulated under different autophagy inducing conditions.

---

There are two major pathways for eukaryotic cells to recycle proteins, lysosomal targeting and the ubiquitin proteasome system (UPS). Within the UPS, an E3 ubiquitin ligase recognizes substrates destined for degradation. In concert with a ubiquitin conjugating enzyme, lysines of the substrate are labeled with ubiquitin or a chain of ubiquitins. The ubiquitinated substrates are recognized by the proteasome and subsequently unfolded and hydrolyzed into short peptides. The peptides are further processed by peptidases into amino acids that become intermediates for various metabolic processes (1, 2). Lysosomes receive substrates either from endocytosis (extracellular and plasma membrane components) or through autophagy (intracellular components). Autophagy, in this paper referring to macroautophagy, utilizes a process where substrates are engulfed into a double membrane compartment known as the autophagosome (3–5). The targeting of substrates to autophagosomes, like the UPS, is often initiated by ubiquitination of target proteins. Ubiquitinated proteins are recognized by autophagy adapters, such as p62/SQSTM1 in humans and Cue5 in yeast, that bind LC3 (Atg8 in yeast) found on expanding pre-autophagosomes (6–9). LC3 binding is facilited by LC3 interacting regions (LIRs and Atg8 interacting motifs, AIMs, in yeast) in autophagy adapters. Closure of the pre-autophagosome captures the substrates. The formed autophagosome then fuses with lysosomes (or vacuoles in yeast and plants) and the content is degraded. Specific transporters export generated amino acids to the cytosol (10). The ability of both the UPS and lysosomal degradation to contribute to the amino acid pool in cells becomes particularly important during physiological conditions that reduce amino acid levels. For example, under conditions of nitrogen starvation, autophagy becomes essential and is the major route for amino acid recycling (11, 12).

Autophagy and the UPS both replenish the amino acid pool as well as utilize ubiquitination as a signal for degradation. Therefore, it makes sense that these processes are coordinated and have some level of redundancy. Indeed, the upregulation of one pathway can, to some extent, compensate for the impairment of the other (13–15). For example, proteasome inhibitors have been shown to induce an autophagic response, and mTOR inhibition by rapamycin, which induces autophagy, was shown to relieve proteotoxic stress caused by proteasome inhibition (16, 17). Further, an increase in proteasome activity was reported in yeast and mammalian cells deleted for the essential autophagy gene *ATG5* as well as upon chemical inhibition of autophagy (18). Similarly, *atg32Δ* cells upregulate proteasome activity in an effort to compensate for an inability to induce non-selective autophagy upon antimycin A treatment (18). Some functions are however clearly unique to each pathway. Proteasomes are, in large, responsible for the degradation of short-lived proteins like IкB (during NFкB signaling) and cyclins (during cell cycle progression) (19, 20). Autophagy, on the other hand, can dispose of aggregated proteins or defective (parts of) organelles directly through the extension of autophagic membranes.

Autophagy is induced by a number of cellular stresses such as nutrient starvation, mitochondrial dysfunction, and various infections (3, 10, 21). This process can capture cytosolic material non-specifically, as well as be selective for specific cargo. Examples of the latter are mitophagy, pexophagy, and ER-phagy, where unique receptors (Atg32, Pex14, and Atg39/40 respectively) ensure specific and efficient targeting to autophagosomes (4, 22– 24). Besides organelles, selective degradation has also been observed for ribosomes in yeast and mammalian cells, a process named ribophagy (24–27) Proteasomes are another multisubunit complex that can be targeted for autophagic degradation, a process referred to as proteaphagy. In yeast, nitrogen starvation and proteasome inhibitor treatment induce proteaphagy (28, 29). The process is conserved as it has also been observed in *Arabidopsis* and human cells (30–32) and appears to be selective. First, receptors have been identified that target proteasomes to autophagosomes: For example, p62 upon amino acid starvation in mammalian cells, (30) and Rpn10 and Cue5 upon proteasome inhibition in plants and yeast respectively (29, 31). Furthermore, several factors that are dispensable for bulk autophagy are important for proteasome autophagy upon nitrogen starvation in yeast (28, 31, 33). How this selective proteasome autophagy is regulated, however, remains poorly understood.

The target of rapamycin complex 1 (TORC1) is a master regulator controlling cell growth and metabolic activity based on the cell’s physiological state. Under nutrient rich conditions, TORC1 is active and general autophagy is inhibited through the phosphorylation and inactivation of Atg13 and Atg1/Ulk1 (34–36). Interestingly, treatment of cells with the TORC1 inhibitor rapamycin has been shown to increase proteasome levels in yeast and mammalian cells (34, 37, 38). Further, an increase in K48-linked ubiquitinated substrates has been observed, along with an increase in proteasomal proteolysis in mammalian cells treated with mTOR inhibitors. Apparently, the capacity of protein degradation by proteasomes becomes important under conditions where TORC1 is not active. Surprisingly, proteasomes undergo autophagic degradation in yeast, plants, and mammalian cells upon starvation conditions well-known to cause TORC1 inhibition (28, 30, 31). To better understand the response of proteasomes to autophagic stimuli, we sought to determine how proteasome autophagy was regulated in yeast. We used various chemical treatments and physiological conditions known to induce general autophagy. Our data show a biphasic response upon inhibition of TORC1 with rapamycin. During the first four hours of treatment, proteasome levels and activity increased. After this, proteasomes were targeted for vacuolar degradation. However, nitrogen starvation, which induces TORC1 inhibition naturally, did not induce a similar response. Indeed, various stimuli that cause TORC1 inhibition and induce general autophagy, showed little to no proteasome autophagy. Thus, proteasome autophagy is regulated distinctly from general autophagy. Consistent with this, we identified several genes required for proteasome autophagy, namely, those encoding the regulatory kinases Mpk1, Mkk1 and Mkk2 as well as the selective autophagy receptor Atg11.

## RESULTS

### Rapamycin induces a bi-phasic proteasome response

Upon nitrogen starvation, proteasomes are targeted for vacuolar degradation via an autophagy dependent pathway. The extent of this degradation appears to vary (28, 31, 33). To elucidate signaling pathways involved, we sought to determine the role of TORC1 inhibition in this process as TORC1 inhibition occurs upon nitrogen starvation and induces autophagy. However, the TORC1 inhibitor rapamycin has been reported to increase proteasome levels and activity (38, 39). To study this contradiction, we monitored the response of proteasomes to rapamycin treatment over an extended time by native gel analyses of yeast lysates. We introduced the sequence of eGFP before the stop codon at the endogenous locus of the core particle (CP) subunit α1 to ensure that α1 transcription is regulated as in wild type and no untagged subunits are produced (40). Upon rapamycin treatment, these cells showed an upregulation of 26S proteasome levels as determined by α1-GFP signal detected on native gel (Fig. 1A, top panel). Coinciding with the increased levels of GFP, we observed more hydrolysis of the suc-LLVY-AMC fluorogenic substrate (Fig. 1A, lower panel). This increase was transient as levels peaked at 30 to 60 minutes. The reduction after the peak was accompanied by an increase in a faster migrating GFP species on the fluorescent scan of the native gel. The accumulation of this species showed kinetics similar to the accumulation of a 25 kDa GFP positive band observed on immunoblots (Fig. 1B) and migrated where we anticipate free GFP to migrate on native gel (41). This “free GFP” is generally indicative of vacuolar targeting (28, 42). We observed the same dynamics when the regulatory particle (RP) subunit Rpn1 was tagged with GFP (Supplementary Fig. 1). Growth for 24 hours in YPD did not yield such an increase in this GFP species and resulted in little to no free GFP on immunoblots (Supplementary Fig. 2). In sum, rapamycin treatment of *Saccharomyces cerevisiae* resulted in an initial upregulation of proteasomes followed by vacuolar degradation.

**Figure 1.**
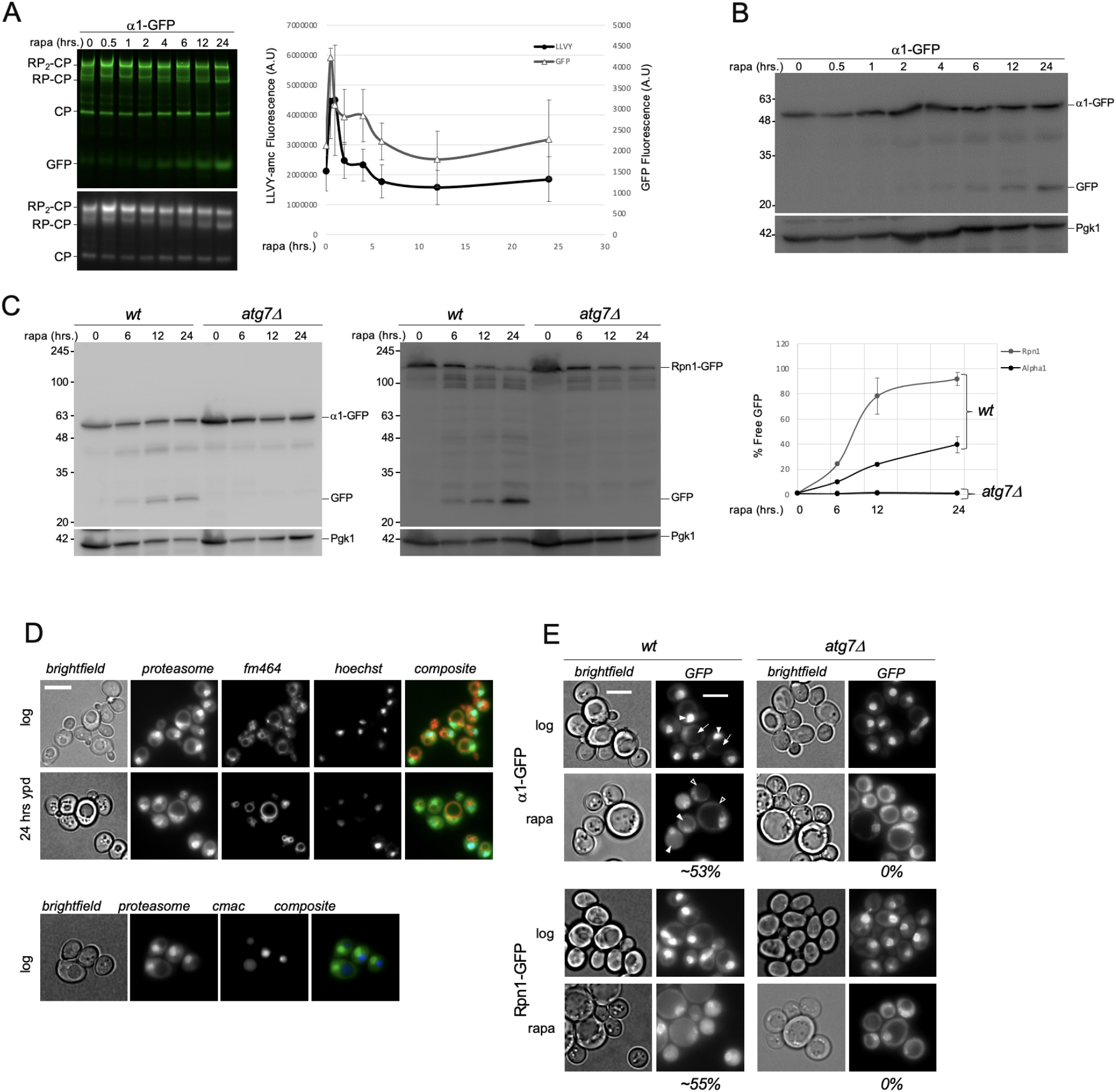
Rapamycin induces a bi-phasic proteasome response. **(A)** Yeast expressing α1-GFP were grown in YPD medium and treated with 200 nM rapamycin. Equal amounts of lysates were separated on native gels and imaged for GFP (top panel) or peptidase activity in the presence of 0.02 % SDS using the proteasome substrate suc-LLVY-AMC (lower panel). Graphs show the quantifications of the levels of GFP and LLVY peptidase activity corresponding to the RP_2_-CP species on native gel, both normalized using PGK1 signal (see (B). Error bars represent SEM. **(B)** Samples from Fig. 1A were denatured and separated on SDS-PAGE and immunoblotted for GFP. Upper band shows α1-GFP and lower band “free” GFP resulting from α1-GFP cleavage by vacuolar hydrolysis. Pgk1 was used as a loading control. **(C)** Wild type and *atg7*Δ yeast expressing α1-GFP or Rpn1-GFP, were inoculated in YPD medium at OD_600_ of 0.5 and grown to log phase (~ 4 hrs). Cells were treated with rapamycin as in (A) and lysed at indicated time points. α1-GFP, Rpn1-GFP, and cleaved free GFP (indicative of vacuolar targeting) were monitored by immunoblotting for GFP. Pgk1 was used as a loading control. Graph shows quantification of immunoblots. **(D)** Rpn1-GFP expressing yeast were stained for the vacuole membrane using FM464 and the nucleus using Hoechst 33342. Microscopy was performed at log phase and following 24hrs growth in rich media (YPD) (top). Rpn1-GFP expressing yeast growing logarithmically were incubated with the vacuole lumen marker CMAC-arg (bottom). Scale bars represent 0.5μm **(E)** Microscopic analysis of yeast collected at log phase and 24 hours after rapamycin treatment. In top image, arrow heads point to nuclei and arrows to vacuoles. In rapamycin treated cells, filled arrow heads indicate cells with vacuolar fluorescent signal, while open arrow heads show subset of non-responding cells. Scale bars represent 0.5μm. Values indicate the percentage of cells in WT and *atg7Δ* with vacuolar GFP signal following rapamycin treatment. Vacuolar GFP signal was rarely (<1 %) observed in non-treated cells. N>100 from 3 biological replicates.

**Figure 2.**
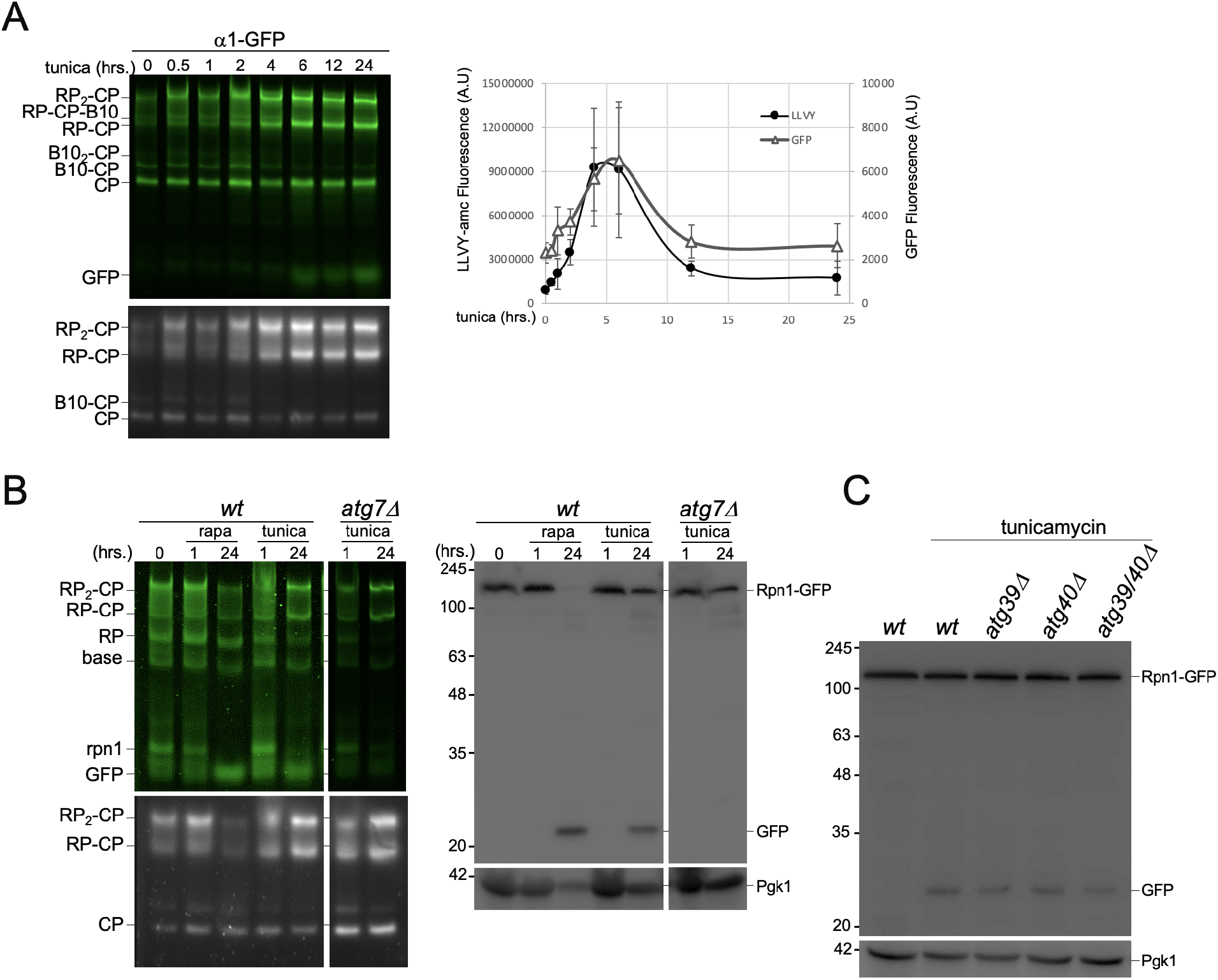
Tunicamycin induces a bi-phasic proteasome response. **(A)** Yeast expressing α1-GFP were treated with tunicamycin (6 μM) for indicated time periods. Cells were collected, lysed under native conditions and equal amounts of lysates were loaded on native gel. Following separation, gels were imaged for GFP fluoresence (top) or suc-LLVY-AMC peptidase activity in the presence of 0.02% SDS (bottom). Quantifications show the amount of GFP signal and LLVY peptidase activity associated with the RP_2_-CP proteasome complexes. GFP and LLVY activity were normalized to Pgk1 intensity using SDS-PAGE immunoblots of the same samples. **(B)** Wild type and *atg7*Δ yeast were treated with tunicamycin and lysates analyzed as in A (left) and denatured for western blotting (right). **(C)** *WT, atg39Δ, atg40Δ*, and *atg39Δ atg40Δ* yeast expressing Rpn1-GFP were treated with tunicamycin as in A. 2 ODs of cells were lysed using the alkaline lysis method. Samples were separated on SDS-PAGE and immunoblotting for GFP and Pgk1 was carried out as described above.

To test if the vacuolar targeting upon rapamycin treatment was autophagy dependent, we introduced Rpn1-GFP or α1-GFP in the autophagy deficient *atg7Δ* strain. No accumulation of “free GFP” was detected in these *atg7Δ* cells indicating that both the proteasome RP and CP, are autophagy substrates upon inhibition of TORC1 (Fig. 1C). Fluorescent microscopy analyses showed that proteasomes are enriched in the nucleus during logarithmic growth (43, 44) with the vacuoles devoid of fluorescent signal. This localization remained similar when cells were grown in rich media for 24 hours (Fig. 1D). While the vacuoles can often be identified with brightfield images, we also confirmed their assignment by staining with FM4-64 (stains vacuolar membrane) or CMAC-Arg (stains vacuolar lumen) (Fig. 1D). Consistent with the gel analyses, 24 hours post rapamycin addition, vacuolar flurescence was observed (Fig. 1E). 55% of wild type cells showed vacuolar localization of proteasomes following rapamycin treatment (n>100, SEM= 1.3 from 3 biological replicates), suggesting this response is less robust than nitrogen starvation, where we observed 90% of the cells with vacuolar fluorescence (28). Consistent with immunoblots, the lack of vacuolar fluorescence in *atg7Δ* cells indicates the translocation is autophagy dependent following rapamycin treatment. Overall, our data show that upon rapamycin treatment, proteasomes undergo a bi-phasic response where an initial upregulation in proteasome levels and activity is followed by an overall reduction via autophagy. Notably, a fraction of generally larger cells showed little proteasome autophagy with rapamycin (Fig. 1E open arrow heads α1 + rapa).

### Proteasome autophagy is distinct from general autophagy

Nitrogen starvation and rapamycin treatment both lead to the inactivation of TORC1 with various downstream effects, such as nuclear translocation of the nitrogen responsive transcription factor Gln3 (34, 45). Thus, rapamycin and nitrogen starvation elicit, at least in part, a similar response through TORC1 inhibition. However, we observed proteasome upregulation only with rapamycin and not upon nitrogen starvation (Fig. 1) (41). This suggests that proteasomes respond differently to these two conditions that induce general autophagy. Therefore, we followed the proteasome response to other conditions and drugs that induce general autophagy.

Tunicamycin is a drug that induces ER stress, the unfolded protein response (UPR), general autophagy, and has been shown to cause proteasome upregulation (38, 46). Cells treated with tunicamycin showed a bi-phasic response similar to rapamycin, however proteasome levels peaked at 4 to 6 hours instead of 1 hour post treatment (Fig. 2A). The increase in free GFP reflects the induction of proteasome autophagy. However, less proteasome autophagy occurred compared to rapamycin treatment as native gels and immunoblots showed a higher ratio of Rpn1-GFP to free GFP and more proteasome peptidase activity was detected 24 hours post tunicamycin addition (Fig. 2B). The accumulation of free GFP was dependent on Atg7, confirming that tunicamycin induced autophagy of proteasomes (Fig. 2B). Since tunicamycin induces both ER autophagy and general autophagy (47), we wondered if the autophagy of proteasomes we observed resulted from ER-associated proteasomes that traffic to the vacuole via ER-phagy. To test this, we deleted ATG39 and ATG40, two genes required for ER/nucleophagy in yeast (23). Neither gene product was required for proteasome autophagy under conditions of nitrogen starvation (29, 33) and we observed no reduction in the amount of free GFP generated upon tunicamycin treatment when ATG39, ATG40 or both genes were deleted (Fig. 2C). Thus, proteasome autophagy observed upon tunicamycin treatment is distinct from ER-phagy.

We next tested another inhibitor of TORC1, caffeine (48, 49). We did not detect an upregulation of proteasome levels or activity when caffeine was added to growing cells, (Supplementary Fig. 3). This difference between caffeine and rapamycin could result from differential inhibition of TORC1 downstream pathways with these drugs, or TORC1-independent effects of rapamycin. However, rapamycin is potent in yeast and both drugs show a similar response in transcriptional profile (50, 51). Consistent with this, proteasome autophagy was also detected following caffeine treatment. However, free GFP accumulated slower and to a lesser extent compared with rapamycin treatment or nitrogen starvation, perhaps because caffeine is less potent as an inhibitor of TORC1 (48, 49).

**Figure 3.**
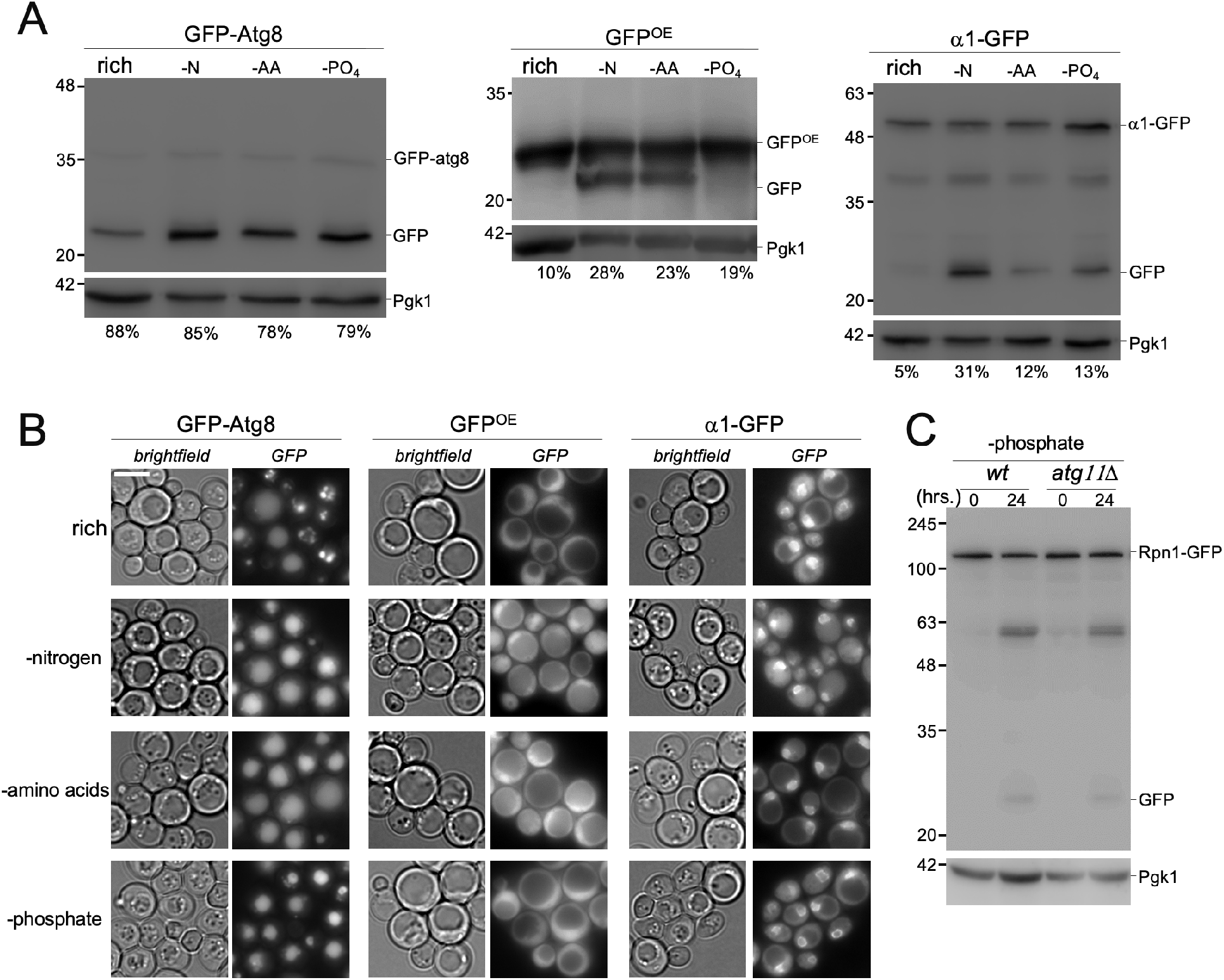
Proteasome autophagy is distinct from general autophagy. **(A)** Yeast strains expressing GFP-atg8, GFP, or α1-GFP, were grown in YPD and subsequently starved 24 hours for nitrogen (-N), amino acids (-AA), or phosphate (-PO_4_). The equivalent of 50 ODs of cells was lysed by cryogrinding. Equal volumes of lysate were blotted for GFP and Pgk1. Value below immunoblots indicate the free GFP signal as percentage of free + fused GFP normalized to PGK1. **(B)** Localization of fluorescent proteins in starved cells from A was monitored by microscopy. Scale bar represents 5μm. **(C)** *WT* and *atg11Δ* yeast were grown to log phase in rich medium and switched to SD medium lacking phosphate. 2 ODs of cells were harvested at indicated timepoints and analyzed as above

Treating *S. cerevisiae* with the different general autophagy inducing drugs, as shown above, led to proteasome autophagy, although each drug elicited different responses with regard to proteasome levels and activity over time. Starving cells for different nutients such as nitrogen, phosphate, amino acids, or carbon, also induces general autophagy (52– 55). To compare general autophagy with proteasome autophagy under these conditions, we utilized GFP-Atg8, which has been used to monitor bulk autophagy (56, 57). While we detected GFP cleaved from Atg8 (indicative of vacuolar targeting) under starvation conditions, we also observed cleavage when cells were grown in rich media for the same amount of time, a condition where little to no bulk autophagy has been reported (Fig. 3A). Considering Atg8 is not only involved in bulk autophagy, but also a number of selective autophagy pathways such as the cytoplasm-to-vacuole targeting pathway (56) this read-out might not be ideal. Therefore, we utilized as a second reporter overexpressed GFP (GFP^OE^) not linked to any other protein. GFP^OE^ does not associate with structures or complexes and its lysosomal processing thus better reflects bulk autophagy. Indeed, when grown in rich media, this reporter did not produce any significant amount of free GFP (Fig. 3A). When we starved cells for amino acids, we observed increased free GFP from GFP-Atg8 compared to rich medium and a similar increase as seen with nitrogen starvation (Fig. 3A and B). This indicates comparable autophagic flux for Atg8 under these conditions, consistent with previous work (53). Free GFP formation in our GFP^OE^ strain was also observed upon amino acid or nitrogen starvation. Proteasome autophagy, on the other hand, was not induced to a similar extent following amino acid starvation compared to nitrogen starvation, as indicated by the reduction in generated free GFP (Fig. 3A).

Phosphate starvation induces general autophagy, albeit to a lesser extent than nitrogen starvation (54). Consistent with this, we found that there was only a modest level of autophagy for GFP^OE^ (Fig. 3A) even though GFP-Atg8 was targeted to vacuoles robustly. This suggests there is little general autophagy induced by phosphate starvation in our strain and the GFP-Atg8 processing we observed could be derived mainly from selective autophagy. This is supported by the reported requirement for ATG11 in phosphate starvation induced autophagy (54) as Atg11 is a protein normally involved in selective autophagy. Proteasomes also appeared to undergo phosphate starvation induced autophagy, albeit to a lesser extent than under nitrogen starvation (Fig. 3A). Atg11 was not required for proteasome autophagy following phosphate starvation, as we did not detect a reduction in the amount of free GFP for the RP upon ATG11 deletion (Fig. 3C).

### Proteasome autophagy requires factors not involved in general autophagy

The conditions utilized above induced general autophagy, but proteasomes were not always robustly targeted for degradation. This suggests proteasomes are not subject to general autophagy but are regulated by a selective process. On the other hand, considering the majority of proteasomes are nuclear, our results could also reflect a lag time for the targeting of nuclear proteins to general autophagy due to the need for nuclear export (Fig. 1D) (43, 44). A hallmark that distinguishes general autophagy from selective autophagy is that the latter pathway depends on specific receptors that recognize and sequester substrates into autophagosomes. For example, selective autophagy of peroxisomes utilizes the receptor Atg30, ER-phagy utilizes Atg39 and/or Atg40, and mitophagy Atg32 or Atg33 (10, 23). All of these factors can interact with the selective autophagy scaffolding factor Atg11 (58, 59). General autophagy, on the other hand, does not depend on Atg11 (60). We have previously reported that nitrogen starvation induced autophagy of an Rpn1-GFP fusion strain was not entirely abolished by a deletion of *ATG17* (28). Interestingly, the residual autophagy of proteasomes in the *atg17Δ* strain was abolished following deletion of *ATG11* (Fig. 4A). For rapamycin induced proteasome autophagy, we were unable to detect free GFP in the *atg17Δ* strain and no difference in free GFP was detected in the *atg11Δ* strain. This suggests that Atg11 is not essential for any form of proteasome autophagy induced by rapamycin. However, less proteasome autophagy is induced with this drug in general, and the levels might be below our detection limit. Other factors required for proteasome autophagy, such as p62 in humans, and Cue5 and Atg24 in yeast, (30, 31, 33), further support the notion that some forms of proteaphagy proceed through a selective pathway.

**Figure 4.**
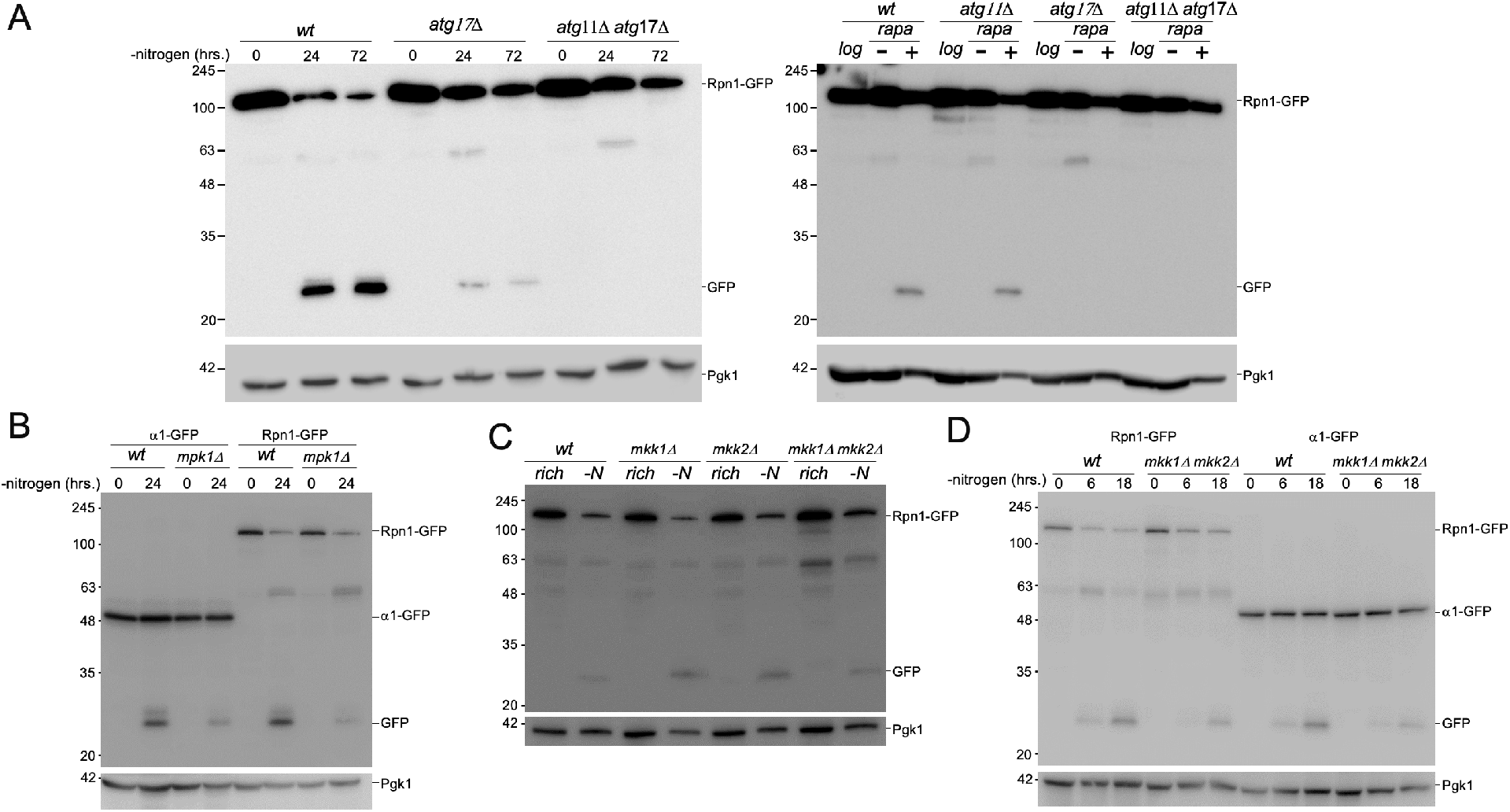
Proteasome autophagy requires factors dispensible for general autophagy. **(A)** *WT, atg11Δ, atg17Δ, atg11Δ atg17Δ* yeast expressing Rpn1-GFP were starved for nitrogen or treated with rapamycin for 24hrs. **(B)** *WT* and *mpk1Δ* cells expressing Rpn1-GFP or α1-GFP were starved for nitrogen and 2 ODs were harvested at indicated time points. Immunoblotting for GFP and Pgk1 were performed as described above. We observed an ~35% reduction in cleaved GFP from Rpn1 and ~33% from α1 (p value 0.067 and 0.0005 respectivey) in the mutants compared to WT 24 hours following starvation. **(C)** *WT, mkk1Δ, mkk2Δ* and *mkk1Δ mkk2Δ* cells expressing Rpn1-GFP were starved for nitrogen and 2 ODs harvested at 24 hours. Immunoblotting for GFP and Pgk1 were performed as described above. **(D)** *WT* and *mkk1Δ mkk2Δ* cells expressing Rpn1-GFP or α1-GFP were starved for nitrogen and 2 ODs harvested at indicated time points. We observed an ~39% reduction in cleaved GFP from Rpn1 and an ~40% reduction from α1 (p=0.030 and 0.119 respectively)

To identify other factors involved in proteasome autophagy, we focused on a key MAP kinase signaling pathway in yeast which utilizes the kinase Mpk1. Mpk1 is not involved in general autophagy (61) but regulates proteasome abundance upon rapamycin addition (38). Deletion of MPK1 in our Rpn1 GFP-tagged proteasome strain resulted in ~35% reduction in the amount of cleaved GFP (going from 62% in wild type to 40% in *mpk1*Δ p=0.067). When α1 was tagged with GFP, we observed a reduction of ~33% (going from 40% in the wild type to 27% in the knock out; p=0.0005) (Fig. 4B). This indicates a role for Mpk1 in efficient proteaphagy upon nitrogen starvation. To trace the kinase pathway involved in this process, we next examined the two kinases directly upstream of Mpk1 in the cell wall integrity pathway, Mkk1 and Mkk2. Mkk1 and Mkk2 are redundant in function and strains deleted of either MKK1 or MKK2 displayed normal proteasome autophagy (Fig. 4C). However, a significant reduction in proteasome autophagy was observed in the MKK1 and MKK2 double deletion mutant upon nitrogen starvation (Fig. 4C, D). For Rpn1, we observed ~39% reduction in cleaved GFP upon deletion of MKK1 and MKK2 (going from 44% in the wild type to 27% in the double knock out; p=0.030). For α1 GFP-tagged strains, the observed reduction in free GFP formed was ~ 40% (going from 25% in the wild type to 15% in *mkk1*Δ *mkk2*Δ strain; p=0.119). These data show that Mkk1 and Mkk2 play redundant roles and, like Mpk1, are important for efficient proteaphagy. Surprisingly, deletion of BCK1, which acts upstream of Mkk1 and Mkk2, did not result in a detectable decrease in proteasome autophagy following nitrogen starvation (Supplementary Fig. 4). This might indicate the signaling does not go through the standard cell wall integrity pathway. Indeed Bck1 has been shown to be dispensible for Mpk1 activation in a number of conditions (62, 63) Nevertheless, the identification of roles for signal transduction (MPK1, MKK1/2) and selective autophagy (Atg11) in proteasome autophagy reinforces the model that proteasomes are specifically targeted for vacuolar degradation.

## DISCUSSION

As an important regulator of many cellular processes, including the cell cycle, the the UPS fulfills an essential function. Similarly, the ability to degrade proteins via autophagy is crucial during mammalian development (64). However, individual cells, like MEFs, grow and multiply without the need for autophagy (65). Similarly, yeast strains lacking the ability to perform autophagy grow at wild type rates under optimal conditions. Autophagy becomes essential, however, upon exposure to certain stress conditions like nitrogen starvation (66). While this indicates important and distinct functions for the UPS and autophagy, it has become clear in recent years that proteasomal and autophagic protein degradation can be synergistic. Here, one system can, to a certain extent, compensate for the impairment of the other (15, 18). Interestingly, in the context of this synergy, recent observations show proteasomes themselves are substrates for autophagic degradation (28–31, 33). While such degradation makes sense under conditions where proteasomes are damaged or not functional (e.g. as a result of inhibition), it is less clear why proteasomes are degraded upon nitrogen starvation. During nitrogen starvation, protein degradation through the UPS as well as autophagy could contribute to replenishing amino acid pools that become depleted. One possible explanation is that translation is largely blocked during nitrogen starvation and ribosomes, which form a substantial amount of the cellular protein mass, are targeted for autophagy (24, 67). Thus, less proteasome activity would be required to degrade (e.g. misfolded) proteins. Another possibility is that cells use highly abundant complexes, such as ribosomes and proteasomes, as a pool of resources that can be utilized to survive nitrogen starvation. The need for the ribophagy receptor NUFIP1 in reactivation of TORC1 signaling upon extended arginine starvation of HEK-293 cells indicates such a role for ribophagy (27).

Approximately 70% of proteasomes in yeast are nuclear and direct autophagy of nuclear material is not responsible for vacuolar targeting of proteasomes. Instead, nuclear export is required for their efficient degradation through autophagy (28, 33). Here, we show that MAPK signalling is important for the efficient degradation of proteasomes, suggesting that specific signaling events regulate proteasome autophagy. A master regulator of metabolic signaling is TORC1. This kinase is inhibited under nitrogen starvation, a condition that can be mimicked with the drug rapamycin. Therefore, the observation that proteasomes are upregulated upon TORC1 inhibition was surprising (38). Our extended analyses showed that proteasomes are indeed initially upregulated. However, following upregulation proteasomes are degraded through autophagy. A similar bi-phasic response was also observed with tunicamycin. Since tunicamycin elicits the unfolded protein response, the upregulation of proteasomes seen with this drug seems logical as proteasomes are important for clearance of misfolded proteins (68, 69). Consistent with this, the transcriptional regulator Rpn4, responsible for upregulation of proteasome genes as well as several other genes, is required for cell survival after inducing ER stress with tunicamycin (70). Proteasome upregulation by rapamycin likely reflects a similar cellular response that benefits cell survival under specific conditions. In line with this, the lack of proteasome autophagy upon amino acid starvation in yeast may indicate that proteasomes contribute to replenishment of amino acid pools. Starvation for nitrogen in yeast also leads to depleted amino acid pools (12, 71), thus the ability of proteasomal degradation to replenish amino acids might be important early in nitrogen starvation. Indeed, we observed delayed proteaphagy compared to general autophagy in yeast (starting ~6 hrs versus ~2 hours) (41, 72).

Another possibility for the different dynamics of proteasomes under autophagy inducing stimuli is that proteasome activity early in autophagy induction may facilitate cellular reprogramming and stress responses. In support of this idea, TORC1 inhibition upon nutrient deprivation has recently been shown to lead to the proteasomal degradation of proteins required for DNA replication (73). In addition, degradation of translation and RNA turnover factors by the proteasome, like Pop2, a deadenylase subunit, and Dcp2, a de-capping enzyme, has been reported upon both nitrogen starvation and rapamycin treatment (67). These data support a role for proteasome activity in regulating protein degradation during autophagy. However, other factors related to ribosome function are degraded via autophagy and do not require proteasome activity (67). Thus, it appears that protein degradation after autophagy induction occurs through both the UPS and autophagy pathways, exemplifying their synergy, and facilitate the appropriate cellular re-reprogramming.

Direct evidence for specificity in proteasome autophagy comes from the identification of specific factors that are required for proteasome autophagy but are dispensable for bulk autophagy. Examples are the requirement for Rpn10 in plants, Atg24 and Cue5 in yeast, and p62 in mammalian cells (29–31, 33). In the current study, we present further evidence in support of specific proteasome autophagy upon nitrogen starvation. We have previously reported that a subset of the regulatory particle was still degraded in an Atg17 mutant (28). The autophagic degradation of the remaining proteasomes depended on Atg11. Atg11 is known to be required for the specific recognition of autophagic cargo in processes like mitophagy, pexophagy, and the CVT pathway (58, 59, 74, 75). Whether or not these proteasomes utilize a selective autophagy receptor that binds Atg11 remains to be determined.

In addition to Atg11, we also observed an important role for the MAP kinase Mpk1 in proteasome autophagy. Importantly, Mpk1 is not involved in general autophagy (61). Currently, it is unclear if Mpk1 facilitates proteasome autophagy by directly phosphorylating proteasomes, regulatory factors, or is part of a longer signaling cascade that ultimately leads to proteasome autophagy. Intriguingly, this kinase is also required for proteasome upregulation following rapamycin addition (38). The role of Mpk1 in rapamycin induced proteasome autophagy could not be determined as *mpk1*Δ cells died approximately 4 hours following rapamycin addition, which is before robust proteasome autophagy is detectable. Mpk1 upregulates proteasomes by inducing expression of regulatory particle assembly chaperones including Adc17. Since, we did not detect a role for Adc17 in proteasome autophagy (supplementary Fig. 5), different factors are likely involved in Mpk1’s roles in the process, a subject we are currently exploring.

When mutants defective for autophagy are starved for nitrogen (28, 40) or treated with rapamycin (Fig. 1E), proteasomes remain nuclear. This raises the question of how proteasome nuclear export/import is linked to autophagy flux. When autophagy is blocked by knocking out Atg7, autophagosomes fail to form (22, 76). However, cellular metabolic conditions and signals that initiate autophagy are still present in nitrogen starved or rapamycin treated cells. For example, in mammalian cells, the nuclear ribophagy receptor NUFIP1 still translocated to the cytosol in ATG7 deleted cells following starvation (27). Thus, proteasomes remaining nuclear suggests that the signals that govern proteasome nuclear export are not present when autophagy is blocked. Proteasome nuclear export potentially does not depend on autophagy induction but instead, autophagic flux. Consistent with this, proteasome autophagy is not observed until approximately six hours post induction (Fig. 1B). Alternatively, it is possible that an equilibrium exists for proteasomes between the cytoplasm and the nucleus. Since only cytoplasmic proteasomes appear to be autophagy substrates, proteasome autophagy results in a drop in the cytosolic proteasome levels. This would induce nuclear export to maintain the nuclear-cytosolic equilibrium of proteasomes. As such, autophagy of nuclear proteasomes would be propagated passively. However, given the dependence of proteasome autophagy on map kinase signaling, this alone does not explain why proteasomes remain nuclear since general autophagy continues. One possibility is that the nuclear cytoplasmic equilibrium of proteasomes is maintained through cellular signaling. This suggest that cells might regulate proteasome autophagy, at least for the nuclear population, by controlling nuclear export. This mechanism is different from protection against autophagy via the formation of proteasome storage granules (77). In all, our data support a model where proteasomes are selective cargo for autophagy regulated through MAP kinase signaling indpedently from general autophagy.

### Experimental procedures

#### Yeast strains

All strains and oligos used in this study are reported in supplementary table S1 and S2 respectively. Our background strains are the W303 derived SUB61 (Matα, *lys2-801 leu2-3, 2-112 ura3-52 his3-Δ200 trp1-1)* and SUB62 (MatA, *lys2-801 leu2-3, 2-112 ura3-52 his3-Δ200 trp1-1*) that arose from a dissection of DF5 (78). Standard PCR based procedures were used to delete specific genes from the genome or introduce sequences at the endogenous locus that resulted in the expression of C-terminal fusions of GFP or mCherry (79–81). GFP-atg8 expressing strains were generated by transformation with BS-Ura3-GFP-Atg8, a gift from Zhiping Xie (Addgene plasmid #69194). To create our GFP-over expressor (OE) reporter, we generated constructs to express GFP from a GPD promotor (GFP^OE^). Plasmid pJR763 (see table S3) was digested with Sac1 and Sal1. This produced a linear DNA fragment for targeted integration in the *URA3-TIM9* region of the yeast genome. To identify successful integration of the expression modules in cells, transformants were grown on plates lacking histidine. Integration was confirmed by PCR.

#### Yeast Growth Conditions

Overnight cultures of yeast were diluted to an OD_600_ of 0.5 and grown in YPD medium to an OD_600_ 1.5 (approximately 4 hours.). Cultures were then treated with drugs for indicated time periods and harvested by centrifugation. Rapamycin was used at 200 nM final concentration, caffeine at 10 mM, and tunicamycin at 6 μM. To induce starvation, cultures growing logarithmically in YPD (2% dextrose) were centrifuged, washed with the respective starvation medium, re-inoculated at an OD_600_ of 1.5, and incubated at 30 °C with constant shaking.

#### Protein lysates and electrophoresis

For native gel analyses, 50 OD’s of cells were pelleted, washed in ddH20, and resuspended in 50 μL ddH_2_O. This suspension was frozen dropwise in liquid nitrogen and stored at −80 °C until further processing. Protein lysates were obtained by cryogrinding cell pellets using CryoCooler, mortar, and pestle from OPS Diagnostics (82). After cryogrinding, lysates were resuspended in lysis buffer (50 mM Tris-HCl [pH 7.5], 1 mM ATP, 5 mM MgCl_2_, 1 mM EDTA). For native gel analyses, equal volumes of lysates were loaded and separated by electrophoresis (90 V, 2.5 to 3 hours., 4 °C). Gels were scanned on Typhoon 9410 or Trio imager (excitation at 488 nm and a 526SP emission filter). Gels were then incubated with suc-LLVY-AMC and imaged using a Syngene G-Box (UV excitation, 440 nm band-pass emission filter) to visualize peptidase activity of proteasome complexes. Samples for SDS-PAGE and western analyses were prepared by mixing equal amounts of lysates for each sample with 1/5 volume of 6X SDS sample buffer (10% SDS, 40 % glycerol, 60mM DTT, 345mM Tris-HCl [pH 6.8] 0.005% bromophenol blue). For experiments not involving native analysis, 2 OD’s of cells were collected at indicated timepoint and stored at −80 °C. Lysis was completed using previously established methods (83). Following electrophoreses, samples were transferred to PVDF membranes and immuno-blotted with antibodies against indicated proteins or tags followed by the appropriate horseradish-peroxidase conjugated secondary antibodies. Antibodies used were anti-GFP (1:500; Roche, cat. nr. #11814460001) and anti-Pgk1 (1:5000; Invitrogen, cat. nr. #459250). Horseradish-peroxidase activity was visualized using the Immobilon Forte Western HRP substrate (Millipore) and images were acquired using the G-box imaging system (Syngene) with GeneSnap software. Data shown are representative of consistently observed trends from at least three independent biological replicates.

#### Fluorescence Microscopy

All microscopy was done with live yeast strains where proteasome subunits where fluorescently tagged (Rpn1-GFP, α1-GFP) at their endogenous locus with expression driven by the endogenous promoter. After indicated treatments, approximately 2 ODs of cells were pelleted, washed with PBS, then resuspended in 30 μL of PBS, and 3 μL mounted on 1% soft agar slides as described by E. Muller (84)(https://www.youtube.com/watch?v=ZrZVbFg9NE8). All imaging by fluorescence microscopy was done within 10 minutes after washing to avoid the effects of prolonged incubation on slides. Routinely, yeast vacuoles were assigned based on phase contrast images. In addition, we regularly stained vacuolar membranes with the dye FM4-64 (Invitrogen; microscope settings: excitation filter 555/28 nm, emission filter of 617/73 nm) or stained the vacuolar lumen using the dye CMAC-Arg (Invitrogen; microscope settings: excitation filter 350/50 nm, emission filter of 457/50 nm). To confirm nuclear localization, DNA was stained using Hoechst (Invitrogen; microscope settings: excitation filter 350/50 nm, emission filter of 457/50 nm). Images were acquired at room temperature using a Nikon Eclipse TE2000-S microscope at 600X magnification with a Plan Apo 60x/1.40 objective Equipped with a Retiga R3™ camera (QImaging). Images were collected using Metamorph software (Molecular Devices) and analyzed using ImageJ.

## Data Availability Statements

All data are contained within the manuscript or the Supplementary Information.

## ACKNOWLEDGEMENTS

We thank members of the Roelofs laboratory for helpful discussions and feedback on the manuscript. This work was supported by grants from the National Institutes of Health; National Institute of General Medical Science (K-INBRE program P20GM103418 and R01GM118660 to J.R.). Further support came from SigmaXI to K.A.W. and support from the Johnson Cancer Research Center at Kansas State University to K.A.W., G.V., and A.B..

