## supplementary figures for "Proteasome autophagy is specifically regulated and requires factors dispensible for general autophagy"

**Running Title**: Proteasome autophagy is distinct from general autophagy

**Supplementary Figures**

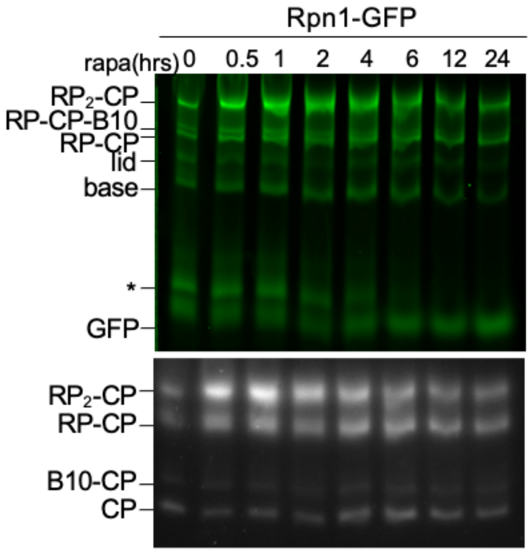


**Supplementary figure 1.** Yeast expressing Rpn1-GFP were treated with rapamycin and samples were collected at the indicated times. Lysates were run on native gel then imaged for GFP and suc-LLVY-AMC peptidase activity.

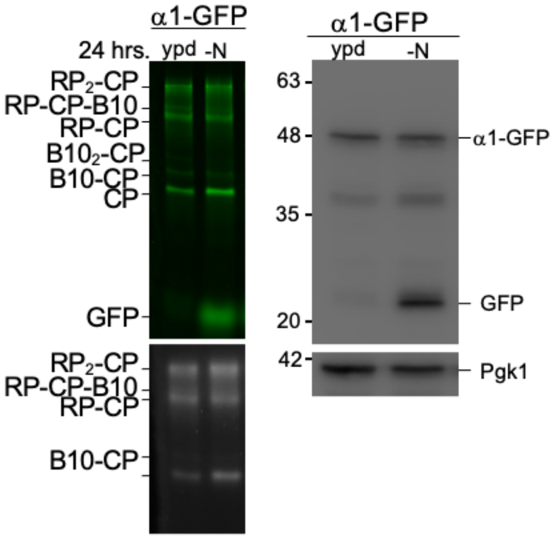


**Supplementary figure 2.** Yeast expressing α1-GFP were grown in YPD media or nitogen starvation media for 24 hours. Samples were collected, lysed, and lysates were seperated on native gel as in supplementary figure 1 or denatured for immuno blotting against GFP or Pgk1.

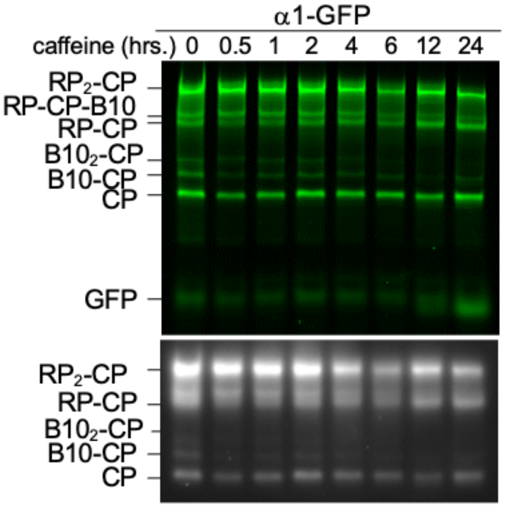


**Supplementary figure 3.** Yeast expressing $\alpha$1-GFP were treated with caffeine, lysed, and analyzed by native gel as in supplementary figure 1.

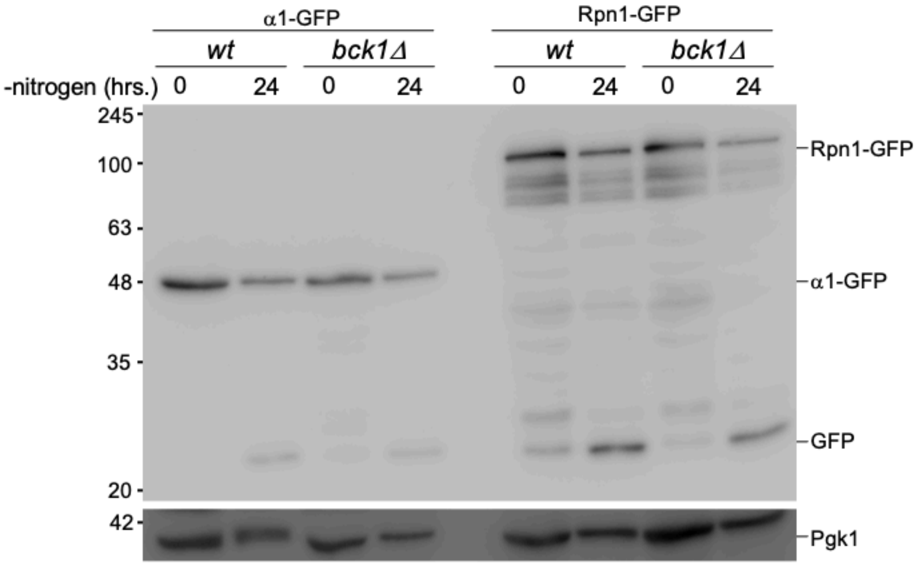


**Supplementary figure 4.** WT and BCK1 deleted yeast expressing $\alpha$1-GFP or Rpn1-GFP were starved for nitrogen and samples were collected at 0 and 24 hours after switching to starvation media. Lysates were separated on SDS-PAGE and immuno blotted for GFP or Pgk1.

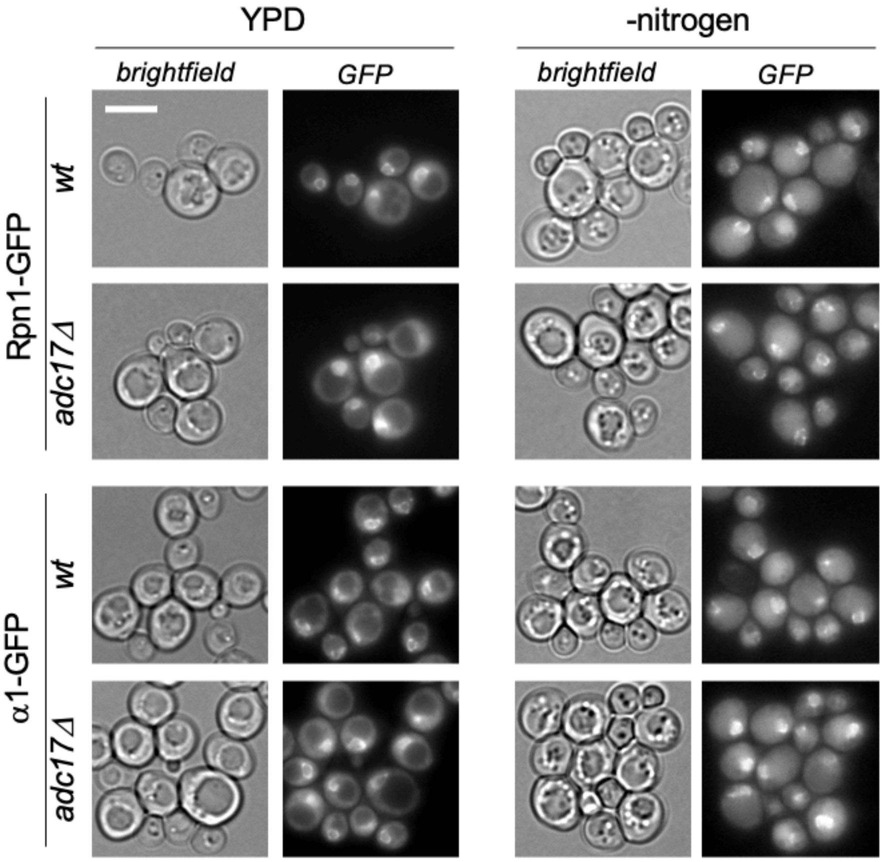


**Supplementary figure 5.** Wildtype and *adc17Δ* yeast expressing Rpn1-GFP or α1-GFP were grown in YPD medium or nitrogen starvation medium for 24hrs and microscopy was performed. Scale bars represent 5 μm.

### Supplementary table 1. strains used in study.

| Strain | Genotype (*lys2-801 leu2-3, 2-112 ura3-52 his3-□200 trp1-1)* | Figure | Source |
|---|---|---|---|
| sJR786 | MAT*α atg7::CloNat rpn1::RPN1-GFP (HIS3)* | **1c,e 2b** | (1) |
| sJR861 | MATα *rpn1::RPN1-GFP (HIS3)* | **1c,d,e 2b,c 3c 4a,b,c,d S1,4** | (1) |
| sJR882 | MATα *rpn1::RPN1-GFP (HIS3) atg17::HYG* | **3d 4a** | (1) |
| sJR900 | MATα *rpn1::RPN1-GFP (HIS3) atg11::G418* | **3c 4a** | (1) |
| sJR923 | MATα *rpn1::RPN1-GFP (HIS3) atg39::HYG* | **2c** | (3) |
| sJR992 | MATα ura3 $^{P}$Atg8-*GFP-atg8 (URA3)* | **3a,b** | (3) |
| sJR1035 | MATα *ura3* $^{P}$*GDP- ubi-met-GFP (HIS5)* | **3a,b** | (3) |
| sJR1084 | MATα *scl1::SCL1-GFP (HIS3)* | **1a,b,c,e 2a, 3a,b 4b,d S2,3,4** | (2) |
| sJR1086 | MATA *atg7::CloNat scl1::SCL1-GFP (HIS3)* | **1c,e** | (2) |
| sJR1091 | MATα *rpn1::RPN1-GFP (HIS3) atg11::G418 atg17::Ura* | **4a** | (3) |
| sJR1151 | MATα *rpn1::RPN1-GFP (HIS3) mpk1::G418* | **4b** | (3) |
| sJR1216 | MATα *scl1::SCL1-GFP (HIS3) mpk1::HYG* | **4b** | (3) |
| sJR1243 | MATα *rpn1::RPN1-GFP (HIS3) atg40::G418* | **2c** | (3) |
| sJR1244 | MATα *rpn1::RPN1-GFP (HIS3) atg39::HYG atg40::G418* | **2c** | (3) |
| sJR1308 | MATα *rpn1::RPN1-GFP (HIS3) adc17::G418* | **S5** | (3) |
| sJR1309 | MATα *scl1::SCL1-GFP (HIS3) adc17::G418* | **S5** | (3) |
| sJR1327 | MATα *scl1::SCL1-GFP (HIS3) bck1::G418* | **S4** | (3) |
| sJR1328 | MATα *rpn1::RPN1-GFP (HIS3) bck1::G418* | **S4** | (3) |
| sJR1387 | MATα *rpn1::RPN1-GFP (HIS3) mkk2::HYG* | **4c** | (3) |
| sJR1390 | MATα *rpn1::RPN1-GFP (HIS3) mkk1::G418* | **4c** | (3) |
| sJR1392 | MATα *rpn1::RPN1-GFP (HIS3) mkk2::HYG mkk1::G418* | **4c,d** | (3) |
| sJR1393 | MATα *scl1::SCL1-GFP (HIS3) mkk2::HYG mkk1::G418* | **4d** | (3) |

a) All strains have the DF5 background genotype ( lys2-801 leu2-3, 2-112 ura3-52 his3-**Δ**200 trp1-1)

1. Waite, K.A., De La Mota-Peynado, A., Vontz, G., and Roelofs, J. (2015) JBC M115.699124
2. Waite, K. A., Burris, A., and Roelofs, J. (2020).. *Sci. Rep.* 10.1038/s41598-020-75126-1

3. This study

### Supplementary table 2. Primers used in this study.

| Primer | Genotype | Template | Sequence (5' to 3') |
|---|---|---|---|
| pRL236 | *atg7::CloNAT* | pAG25[1] | TTCATTATATTTCAACAAATATAAGATAATCAAGAATAAACGTAC GCTGCAGGTCGACG |
| pRL237 | *atg7::CloNAT* | pAG25[1] | CGGAAAGTGGCACCACAATATGTACCAATGCTATTATATGCAATC GATGAATTCGAGCTCG |
| pRL293 | *atg17::HYG* | pFA6a-hphNT1[2] | ATTCGATACTGCGAGGATATTATCAACGTATTTAACACCTCGTAC GCTGCAGGTCGAC |
| pRL294 | *atg17::HYG* | pFA6a-hphNT1[2] | GATACAATTATTGAATCTTTGTACCGTATCCTTTTTTTCCTATCGA TGAATTCGAGCTCG |
| pRL340 | *atg11::G418* | pFA6a-hphNT1[2] | GTTGTTCGGAAAGTACTTCTTTTATTTTCTTTTATACATCCGTACG CTGCAGGTCGAC |
| pRL341 | *atg11:: G418* | pFA6a-kanMX6[2] | ACATAATTAAAATCTTGTCATTTGTGACAAACGTTTAGCACATCG ATGAATTCGAGCTCG |
| pRL371 | *atg39::HYG* | pFA6a-kanMX6[2] | TAATAGAGACTAGTAAAACAGTCGAGTTGTCGGACCTAAACGTA CGCTGCAGGTCGAC |
| pRL372 | *atg39::HYG* | pFA6a-hphNT1[2] | CTTTTGTTAATTTCATTCTTCATGCTGGGTTTTGGATGATATCGAT GAATTCGAGCTCG |
| pRL451 | *mpk1::G418* | pFA6a-kanMX6 | GTAGAAATAATTGAAGGGCGTGTATAACAATTCTGGGAGCGTAC GCTGCAGGTCGAC |
| pRL452 | *mpk1::G418* | pFA6a-kanMX6 | GCTTACATCTATGGTGATTCTATACTTCCCCGGTTACTTATAGATC GATGAATTCGAGCTCG |
| pRL690 | *atg40::G418* | pFA6a-kanMX6 | ACGTTCTTTCTGCTGTGCTTCACTCCACCATAGAAAACTACGTAC GCTGCAGGTCGACG |
| pRL691 | *atg40::G418* | pFA6a-kanMX6 | CTTCATAGACTACCATTATGGTAAAATGGAAAAACTATTCATCGA TGAATTCGAGCTCG |
| pRL694 | *bck1::G418* | pFA6a-kanMX6 | CACTAAATATAGTATTAAAATAGTTCAACTCCACCTCCAACGTAC GCTGCAGGTCGACG |
| pRL695 | *bck1::G418* | pFA6a-kanMX6 | CGTATGCATAAATATCTTAAGTATAGATCGATCCTAATAGATCGA TGAATTCGAGCTCG |
| pRL718 | *adc17::G418* | pFA6a-kanMX6 | TAAAGCAAATCAAAACATAATAACTACTACAAGTAACATACGTA CGCTGCAGGTCGACG |
| pRL719 | *adc17::G418* | pFA6a-kanMX6 | TGACGTGAAAATGATGCGCAGTAAAACTAAATCCCGTCTCATCG ATGAATTCGAGCTCG |
| pRL792 | *mkk2::HYG* | pFA6a-hphNT1[2] | GTTATCATATCTACAAAATACCAATTATATACACAGGATACGTAC GCTGCAGGTCGACG |
| pRL793 | *mkk2::HYG* | pFA6a-hphNT1[2] | AAAAAGTCAGTTCTGGTTACGAGAGGAAAATGTTGGAAGTATCG ATGAATTCGAGCTCG |
| pRL796 | *mkk1::G418* | pFA6a-kanMX6 | CCAAATTAACTCTATCCTTCCATTGCACAATTTGCCCAGTCGTAC GCTGCAGGTCGACG |
| pRL797 | *mkk1::G418* | pFA6a-kanMX6 | AAAATAAACTTAATCATGTTCGCAAAAAATTGCTTATTGAATCGA TGAATTCGAGCTCG |


1. Goldstein, A. L. & McCusker, J. H. Three new dominant drug resistance cassettes for gene disruption in Saccharomyces cerevisiae. Yeast 15, 1541–53 (1999).
2. Janke, C. et al. A versatile toolbox for PCR-based tagging of yeast genes: new fluorescent proteins, more markers and promoter substitution cassettes. Yeast 21, 947–62 (2004).
3. Hailey DW, Davis TN, Muller EG. Fluorescence resonance energy transfer using color variants of green fluorescent protein. Methods Enzymol. 351:34-49 (2002.
